# Note on the power law of forgetting

**DOI:** 10.1101/173765

**Authors:** Michael J. Kahana, Mark Adler

## Abstract

Power functions (e.g., *f*(*t*) = *at*^−*b*^) describe the relationships among many variables observed in nature. One example of this is the power law of forgetting: The decline in memory performance with time or intervening events is well fit by a power function. This simple functional relationship accounts for a great deal of accumulated data. In this note, we consider a simple yet general memory model in which all items decay monotonically in strength, but at different rates. To translate between continuous changes in strength and actual memory for events we assume a simple strength threshold for remembering. We prove a limit theorem for this model: as time grows large and memories decay, the empirical forgetting function approaches a power function under very general conditions. Power forgetting emerges for almost any monotonically decreasing strength function (including linear and exponential cases). We also illustrate by way of simulations that the power function provides an excellent fit to the entire time-course of the forgetting function, not just its limiting behavior.

## 1. INTRODUCTION

Ever since Ebbinghaus (1885/1913) inaugurated the scientific study of memory, researchers have examined the manner in which memory performance declines with time or intervening events (i.e., the forgetting function). Although it has long been known that forgetting occurs rapidly at first and more slowly as time goes on, it was not until quite recently that the mathematical form of the forgetting function has been precisely established. Wixted and colleagues (Wixted, 1990; Wixted & Ebbesen, 1991, 1997) have demonstrated that the form of forgetting, across various materials and memory tests, is characterized mathematically by a power function. Rubin & Wenzel (1996) compared over a hundred forgetting functions and found that the power function was one of only four that provided a good fit to a wide range of forgetting data. That is, accuracy in a memory task at time *t* is given by *y* = *at*^−*b*^, where *a* and *b* are positive real numbers. This invariance in the form of forgetting suggests a basic law of human memory.

If the nature of the functions that characterize learning and forgetting can teach us about their underlying processes, we ought to take any evidence either for or against a particular function very seriously. For instance, if a model assumes that memory trace strength decreases in an exponential fashion, a seemingly obvious implication is that such a model should be rejected. How could exponentially decreasing trace strength be reconciled with power-law forgetting?

Exponential strength decay is a natural assumption for memory models. If you consider a model like TODAM (Murdock, 1982, 1997), forgetting arises due to a forgetting parameter, *α*, and also because the variance of the memory increases with list length (but see Murdock & Kahana, 1993,b). In TODAM, the storage equation for item information (Murdock & Lamon, 1988) is given by:

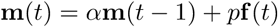

Because the memory is premultiplied by *α* as each new item is learned, recognition performance (as measured by *d′*) should decrease as a function of *α* to the power of lag (or time).

Exponential decay of trace strength or cue-target strength is common in other models as well. Mensink & Raaijmakers (1988) have a forgetting process based on contextual drift that is *nearly* exponential. The same is true of other models of contextual drift (Murdock, 1997; Howard & Kahana, 2002). Bower’s (1967) multicomponent model also assumes exponential strength-decay of the individual components.

Although exponential decay in trace strength is a common assumption of memory models, the data strongly suggest that forgetting obeys a power law. Such power laws are not unique to forgetting. Learning is also well described by a power law. The reduction in reaction times that comes with practice is a power function of the number of training trials (see Anderson, 1995, for a review). Indeed, power laws describe a great many natural phenomena ranging from sensory scaling to the distribution of city sizes. What can we learn from the power-law of forgetting? Does a power-law of forgetting imply that models based on exponential strength decay mechanisms must be rejected?

### 1.1. Interpretive Problems

Using a computational analysis, Anderson and Tweney 1997 showed that arithmetic averaging of exponential functions can give rise to power functions. They also reported that they were unable to find a general analytic solution to the problem of aggregated exponentials. Indeed, this problem may not have a solution without adding some simplifying assumptions. Anderson & Tweney (1997) concluded that the power law of forgetting may be an artifact of arithmetic averaging (see also Anderson & Tweney, 1998)

Responding to the Anderson & Tweney (1997) critique of power functions, Wixted & Ebbesen (1997) showed that even with geometric averaging of individual subjects’ data, a power function still fits the data better than an exponential function. Of course, it is possible that the problem is not at the level of subjects, but rather at the level of items, or even subject-item interactions. Wixted & Ebbesen (1997) acknowledge the possibility that exponential forgetting with variability in forgetting rates across items could give rise to aggregate power functions of forgetting.

Wickens (1998) pursued this point, showing that very different theories of forgetting can give rise to very similar-looking retention functions. In particular, starting with the assumption that individual items are forgotten exponentially, Wickens (1998) showed that models based on (1) heterogeneity of forgetting rates, (2) consolidation of traces over time, and (3) competition among traces for survival, under certain distributional assumptions, can give rise to a Pareto II distribution of item survival. For large *t*, this description of forgetting simplifies to the power law that has been shown to well describe the empirical data.

Although power-functions provide a good fit to forgetting data for ranges of the forgetting function, there is no sense in which a complete forgetting function, with performance measured from *t* = 0 to *t* = ∞, could be well fit by a power function. This is because a power function implies that the measure of performance tends to infinity as *t* tends to zero. With percent correct as the dependent measure, performance at *t* = 0 will tend to 1.0, thus “rejecting” a power function.

In the work presented here, we concentrate on the asympototic properties of forgetting — the behavior of the forgetting function as *t* → ∞. In particular we obtain analytic results that show that ast *t* → ∞, performance is a power function of *t* for a simple but general class of forgetting models. Simulations support these analytic results, and show that the emergence of power forgetting appears quickly for many different parameter values of the underlying models.

### 1.2. Power forgetting in a strength-decay to threshold memory model

Memory researchers are interested in knowing the form of changes in memory strength over time — not changes in memory performance over time. But, we can’t measure memory strength directly; rather, we measure performance. In most memory tasks, performance is based on a summary statistic over discrete events (success or failure in recognizing or recalling an item at a given delay between study and test).

Consider the following very simple, yet general, memory model. Items stored in memory have a strength value, *S*. In the absence of reinforcement but the presence of intervening activity, the strength of each item decays monotonically according to some strength decay function.

Assuming that the strength decays exponentially: *S_i_*(*t*) = *αS_i_*(*t* − 1) and 0 < *α* < 1, therefore,

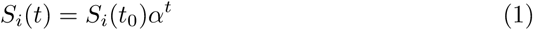

This type of difference equation is characteristic of a number of memory models (e.g., Murdock, 1982, 1997; Mensink & Raaijmakers, 1988). But the form of the forgetting function turns out not to be crucial. The important consideration is that when an item’s strength falls below a fixed threshold, *k*, the item is forgotten. So long as the strength is greater than *k*, the item is remembered. We also must assume that strength decays at different rates for different items (cf. Anderson & Tweney, 1997; Wickens, 1998). The basic features of the model are illustrated for an exponential strength decay function in Figure 1.

**FIG. 1.**
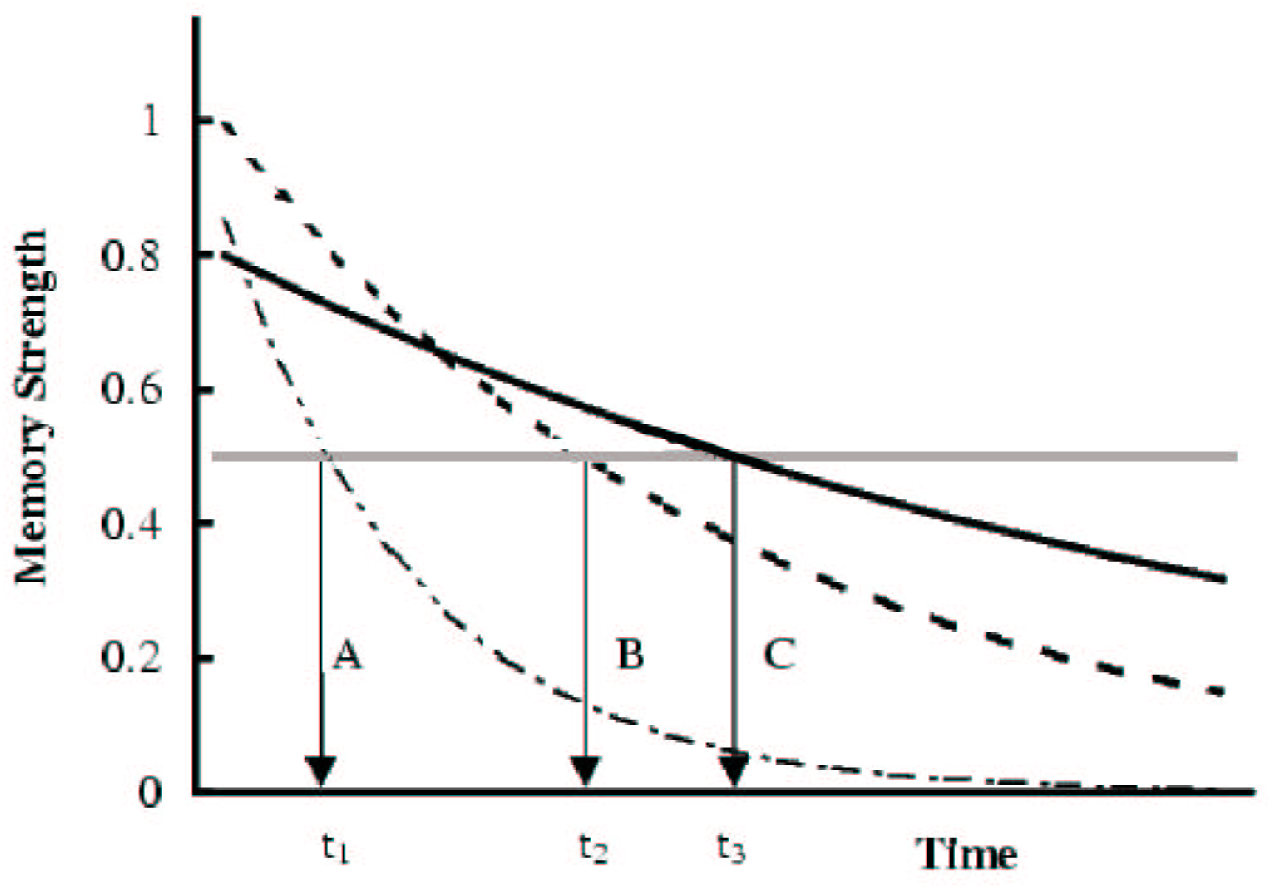
A simple model of recall. Three curves illustrate the reduction in memory strength for three different items (A, B, and C). Items are assumed to decay at different rates and with different initial strengths. The gray bar indicates the strength threshold for correct recall or recognition. Items are forgotten when their strength falls below the threshold. Three arrows indicate the points in time when items A, B, and C are forgotten. Before *t*_1_ all three items are remembered, two items are remembered until *t*_2_, one item is remembered until *t*_3_, and after *t*_3_ none of the three items are remembered.

Here we show that under these conditions, the average forgetting function approaches a power function as *t* grows large. Before proving a limit theorem under fairly general conditions, we present proofs for two special cases: linear trace decay and exponential trace decay. Simulation results for the case of exponential strength decay show that power forgetting emerges quite rapidly and is not just a property of the tail of the forgetting function (i.e., when accuracy approaches zero at very large *t*). In the special cases we assume, for convenience, that strength decay rates are normally distributed subject to the constraint that all items do exhibit at least some decay (we use truncated Gaussians). In the general derivation, we show that this assumption is not at all crucial to the proof of the limit theorem.

### 1.3. Special Case 1: Linear Strength Decay

Consider a linear strength-decay function with independent variable coefficients: *S_i_*(*t*) = *a_i_* − *b_i_t* with *a_i_* ~ *N*(*μ_a_*, *σ_a_*), *b_i_* ~ *N*(*μ_b_*, *σ_b_*), and *μ_a_* > 0, *μ_b_* > 0. Next, define a counter of retrievable memories, *r_i_*(*t*), such that:

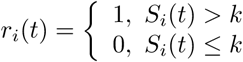

This is a key feature of the model. Items are retrieved if strength is greater than *k*; otherwise, they are forgotten.

Observe that:
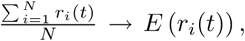
as *N* → ∞ by the Law of Large Numbers, because the *r_i_*(*t*) are identically distributed, independent random variables. The expectation of *r_i_*(*t*), *E*(*r_i_*(*t*)) is our forgetting function, averaged across many discrete items. Computing this expectation will give us the form of the forgetting function under the conditions outlined above. To compute *E*(*r_i_*(*t*) we proceed as follows:

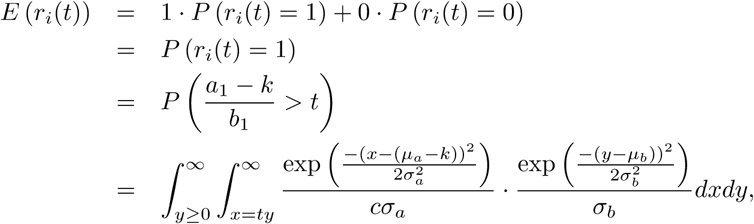

where we let *x* = *a*_1_ − *k* and *y* = *b*_1_, and have constructed Gaussians about these points. Note that we are using a truncated Gaussian about *b_i_*, if *b_i_* ≥ 0, as seen by the lower limit of the outer integral (*y* ≥ 0), and
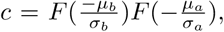
with *F* defined below.

Now we set
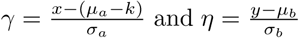
to simplify our computations. To make a change of variables, however, we must modify the limits of integration. We see that
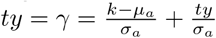
where *y* = *ησ_b_* + *μ_b_*. In which case we can write:

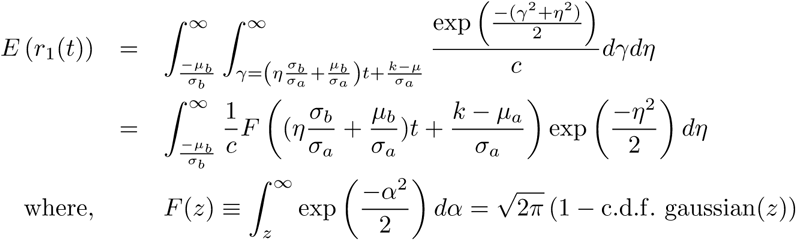

Thus, *F*(−∞) = 1, and
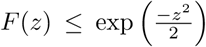
for large *z* > 0, which is easily integrable. Now setting
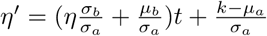
we find that:

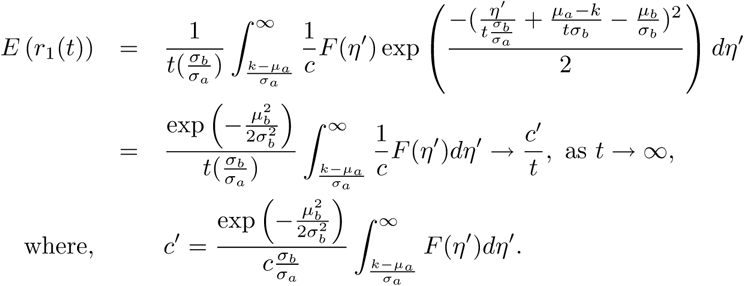

### 1.4. Special Case 2: Exponential Trace Decay

Following the same construction as in the linear case, we define a counter of retrievable memories:

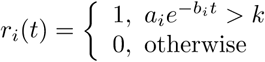

Further, let *x* = *a_i_* and *y* = *b_i_*. Then the probability of retrieving a memory is given by:

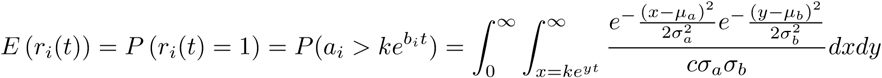

where
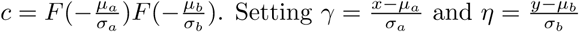
then gives

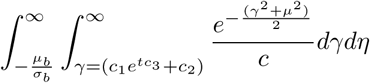

where
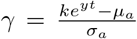
and *y* = *ησ_b_* + *μ_b_*, and where
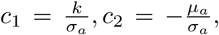
and *c*_3_ = *ησ_b_* + *μ_b_*. We then obtain

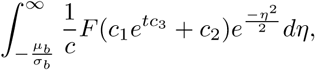

where *F*(·) is as defined previously. Setting *tc*_3_ = *t*(*ησ_b_* + *μ_b_*) = *η*′ gives

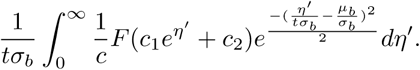

Thus, as *t* → ∞,

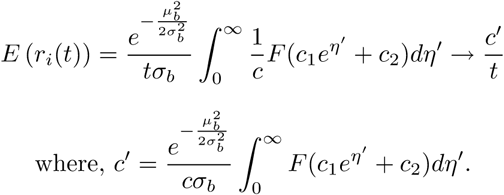

### 1.5. Discussion of the special cases

In the two special cases presented above, we saw that as *t* → ∞ a threshold model in which item strength decays either linearly or exponentially (with time) yields a hyperbolic relationship between recall probability and time. Although this is a special case of the power law of forgetting, the exponents for the limiting case of *t* → ∞ are exactly 1, whereas in actual experiments, the exponent that fits the full range of participants’ performance is often significantly less than one.

To see that the model does actually produce exponents that are less than one consider the following analysis. Recall that the forgetting function, given by *E*(*r_i_*(*t*)), can be rewritten as follows:

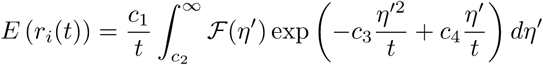

In the equation above, *c*_1_, *c*_2_, *c*_3_ and *c*_4_ are all constants. If we use the analytic series,
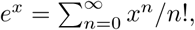
, for *x* ∈ 𝓒, then we can rewrite the above expression as:

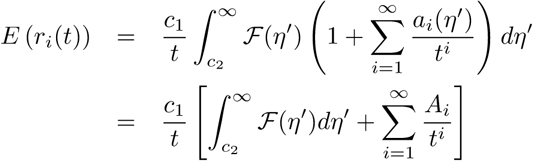

Next, setting *c′* = *c*_1_*A*_0_, and *b_i_* = *A_i_/A*_0_, we have:

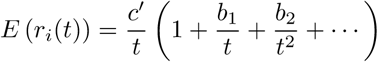

If we define

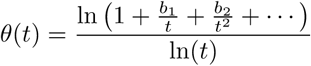

then we have

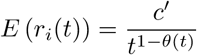

Because *θ*(*t*) < 1 and *θ*(*t*) → 0 as *t* → ∞, we can see how the exponent of the power function varies with *t* in a systematic manner. For smaller values of *t*, the exponent will be less than one, but it will approach 1 as *t* grows large.

The analysis presented above shows that the simple decay-to-threshold model gives rise to power forgetting even when the strength decay functions are linear or exponential. In the next section we consider the centrality of our assumptions in producing the “power-law” behavior at large *t*. In particular, we generalize our analytic results to a very a broad class of strength decay functions and to nonparametric variability in the decay parameters.

### 1.6. General Derivation

Consider a strength-decay function of a general form:
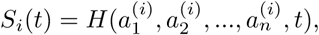
and assume the following conditions:

1. *H* = *H*(*a*_1_, *a*_2_*e*_2_(*t*), *a*_3_*e*_3_(*t*),…,*a_n_e_n_*(*t*)) with

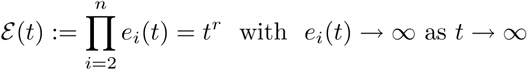 #### Remark 1

This form of the strength decay function is very general. Consider the case of the exponential decay common to many memory models, *S_i_* (*t*) = *ae^−*bt*^.* In this case, *a*_1_ = *a*, *a*_2_ = *b*, and *e*_2_(*t*) = *t*. Similarly, the form of *H* also accommodates linear strength decay.
2. *H* is globally monotonic increasing in its first argument
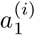
for all values of the other arguments, monotonic decreasing in *t*, and smooth (*l* continuous derivatives) in all arguments.
3. 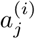
has distribution function
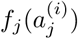
with *f_j_* independent of *i*, just depending on *j*, such that *f_j_* (*x*) = 0 for *x* < 0,
4. all random variables are independent.

#### Remark 2.

Condition 2 implies that
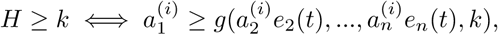
and
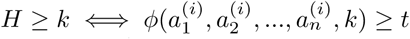
for some smooth function (*l* − 1 continuous derivatives) in all arguments by the global monotonicity and implicit differentiation.

Set
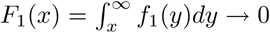
as *x* → ∞. Define a counter of retrievable memories, *r_i_*(*t*), such that:

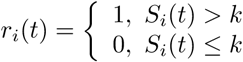

Again, this is a key feature of the model. Items are retrieved if strength is greater than *k*; otherwise, they are forgotten. As before, observe that
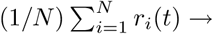
*E* (*r_i_*(*t*)), as *N* → ∞, by the Law of Large Numbers, because the *r_i_* (*t*) are identically distributed, independent random variables.

The expectation of *r_i_*(*t*) is our forgetting function, averaged across many discrete items. Computing this expectation will give us the form of the forgetting function under the general conditions outlined above. To compute *E*(*r_i_*(*t*)) we proceed as follows:

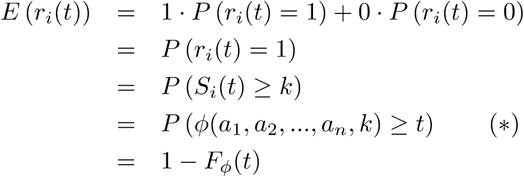

where we have set all
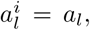
and where *F_ϕ_* is a c.d.f. of the random variable *ϕ*(*a*_1_, *a*_2_,…, *a_n_*). Note that we have used in the above that

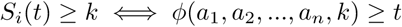

for some function *ϕ* by *Remark 2*.

Analytic solutions for the special cases of linear and exponential forgetting functions (shown earlier) suggested that power forgetting emerges as a general property of this model. Here we approach the general solution by asking the prospective question: When does

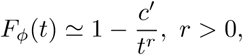

for *t* large? In other words, what are the conditions that would result in power-law forgetting?

This requires that the density distribution function

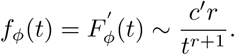

This is really a condition on the function *ϕ*(*a*_1_, *a*_2_, …, *a_n_*, *k*).

Further analysis proceeds as follows: Using (^∗^) and Remark 2 we compute in general that:

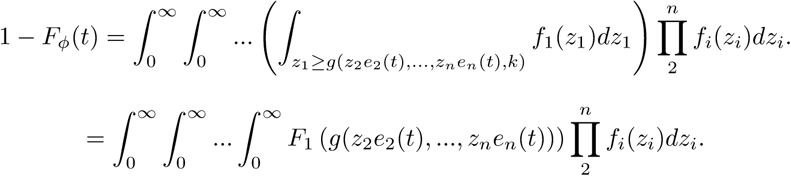

Setting
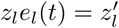
gives

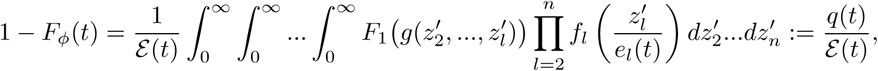

Using *e_l_*(*t*) → ∞ as *t* → ∞ (so as
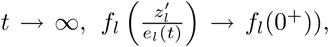
and using Lesbesgue’s dominated convergence theorem and Assumption 3 below, we see that as *t* → ∞, 1 − *F_ϕ_*(*t*) approaches:

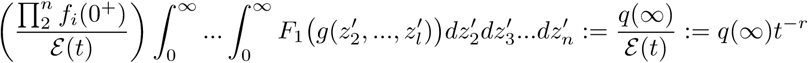

by our hypothesis, with,

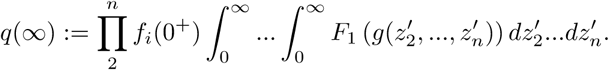

In order to obtain the limit above, we assumed that

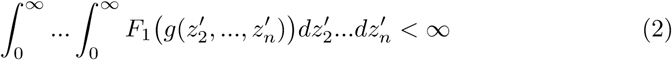

and the mild assumption that there exists an *M* such that |*f_i_*(*x*)| < *M* for all *i* and *x* (so that we can use dominated convergence).

The rate of convergence of *q*(*t*) → *q*(∞) is controlled by the decay rate of *F*_1_(*g*(*z*′)) as *z* → ∞. For very small *t*, it is likely that the strengths of the individual items are all > *k*. This would mean that nothing has been forgotten yet. As *t* grows, and strengths begin to drop below the threshold, *k*, power forgetting may appear quite quickly.

### 1.7. Simulations

To examine how rapidly power forgetting emerges in our model we simulated the special case of exponential trace decay. In particular, *r*(*t*) = *E*(*a_i_e*^−*b_i_t*^ > *k*), where *a* and *b* are random variables. In Simulation 1, *a* and *b* were truncated Gaussians (*a_i_* > 0 and *b_i_* > 0). This follows the proof of the special cases. In Simulations 2 and 3, *a* and *b* were drawn from an exponential distribution. This avoids the problem of having to truncate negative values. In addition to the parameters that determine the distributions of *a* and *b*, the model has one additional parameter: the forgetting threshold, *k*.

Figures 2-4 show three simulation runs with different parameter values. For each simulation we have plotted the models’ predictions in both linear and log-log coordinates (a power function appears as a straight line in a log-log plot). The parameter values and *r*^2^s are given in the figure legends.

**FIG. 2.**
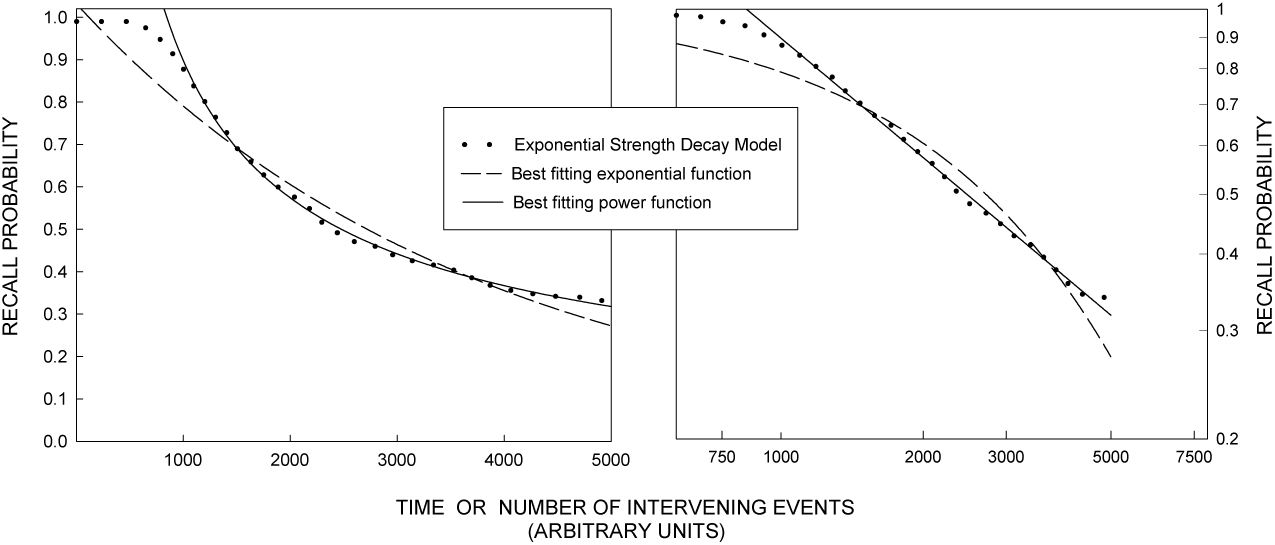
Simulation example #1: Forgetting function for an exponential strength decay model. Left panel plots data in linear coordinates; Right panel plots data in log-log coordinates. The following parameter values were used: *k* = 0.001, *a* ~ *N*(2.5,1.0), *b* ~ *N*(.003, .004). For this simulation, *a* and *b* are truncated normals, with 500 items being forgotten at different rates. Exponential and power functions were fit to data in the range of *t* = 1000 to 5000. The best fitting power function is given by P(recall)= 77*t*^−.64^ with *r*^2^ = 0.999. The best fitting exponential function is given by P(recall)= 1.03*e*^−.0003*t*^ with *r*^2^ = 0.95.

**FIG. 3.**
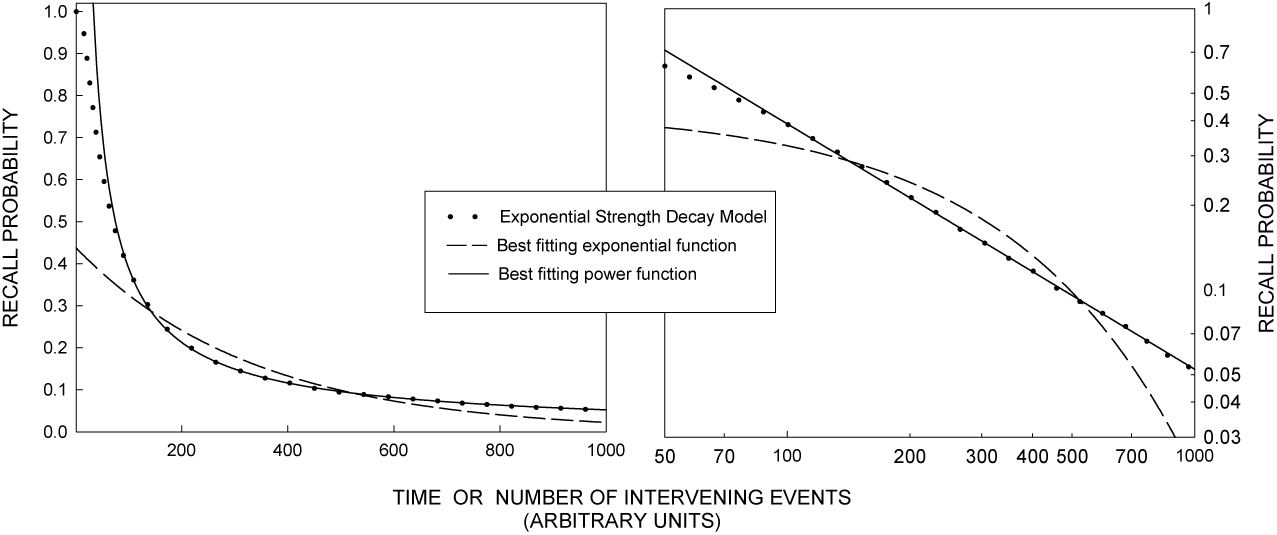
Simulation example #2: Forgetting function for an exponential strength decay model. Left panel plots data in linear coordinates; Right panel plots data in log-log coordinates. The following parameter values were used: *k* = 0.2, *a* ~ *N*(3.5,0.5), *b* ~ *f* (.06) where *f* denotes an exponential distribution. For this simulation, 12,000 items are forgotten at different rates. Exponential and power functions were fit to data in the range of *t* = 80 to 1000. The best fitting power function is given by P(recall)= 21*t*^−.87^ with *r*^2^ = 0.999. The best fitting exponential function is given by P(recall)= 0.44*e*^−.003t^ with *r*^2^ = 0.92. Identical fits were obtained with the following parameter values: *k* = 1, *b* ~ *f* (.025).

**FIG. 4.**
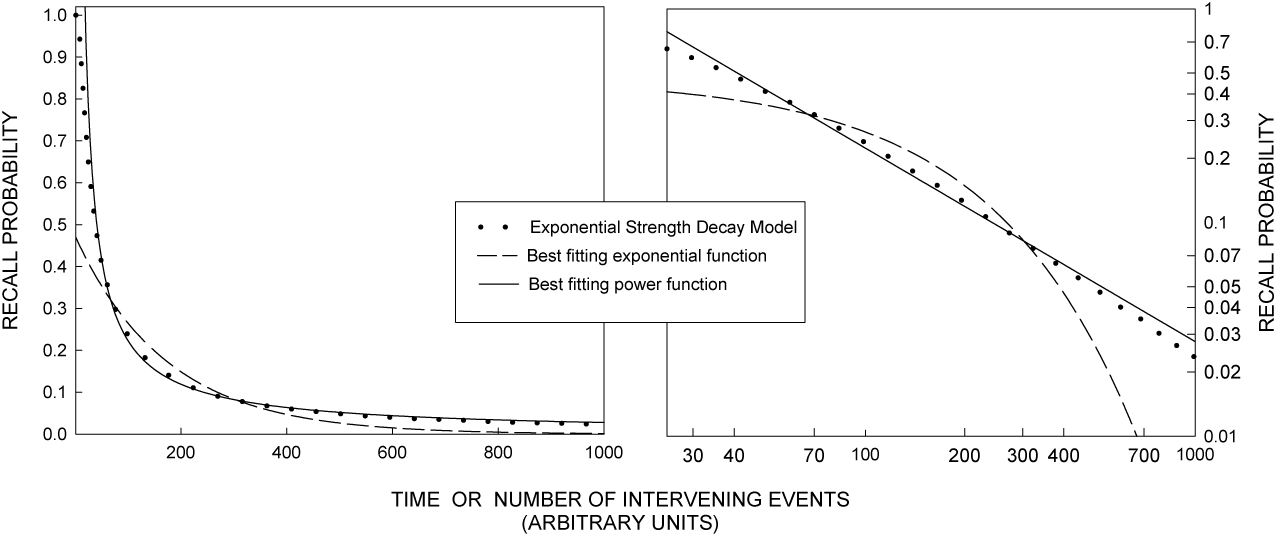
Simulation example #3: Forgetting function for an exponential strength decay model. Left panel plots data in linear coordinates; Right panel plots data in log-log coordinates. The following parameter values were used: *k* = 0.02, *a* ~ *N*(1.0,0.2), *b* ~ *f*(0.06) where f denotes an exponential distribution. For this simulation, 10,000 items are forgotten at different rates. Exponential and power functions were fit to data in the range of *t* = 35 to 1000. The best fitting power function is given by P(recall)= 14.4*t*^−.91^ with *r*^2^ = 0.995. The best fitting exponential function is given by P(recall)= 0.47*e*^−.006*t*^ with *r*^2^ = 0.93. Consistent with the theoretical analysis, the power function that best fit data in the range of *t* = 150 to 1000 was given by P(recall)= 29.3*t*^−.103^ with *r*^2^ = 1.000

These simulations show that a simple decay-to-threshold model with exponentially decaying memories can produce aggregate power functions. These results are typical of many simulations run with other parameter values. Although the power function provides an excellent fit once performance drops significantly below ceiling, the power function does not provide an adequate fit to the early portion of our model’s forgetting functions^1^. Like experimental subjects, our model dictates that performance in the first few moments after learning remains at ceiling. Because *at*^−*b*^ → ∞ as *t* → 0 the power function is unable to fit this aspect of either the human data or the model results. It is only once forgetting is well underway that the power form emerges. This is consistent with our analytic results showing that power forgetting emerges as *t* grows large.

The simulations described above all assumed exponential strength decay of individual memories. Under these assumptions, the exponent of the power function that best fits the data should approach 1 as *t* → ∞. Figure 5 replots the simulation results shown in Figure 4 and shows separate power-function fits to different ranges of the forgetting function. Consistent with the analytic results, the best fitting exponents decrease monotonically to -1 as we fit later regions of the forgetting curve.

**FIG. 5.**
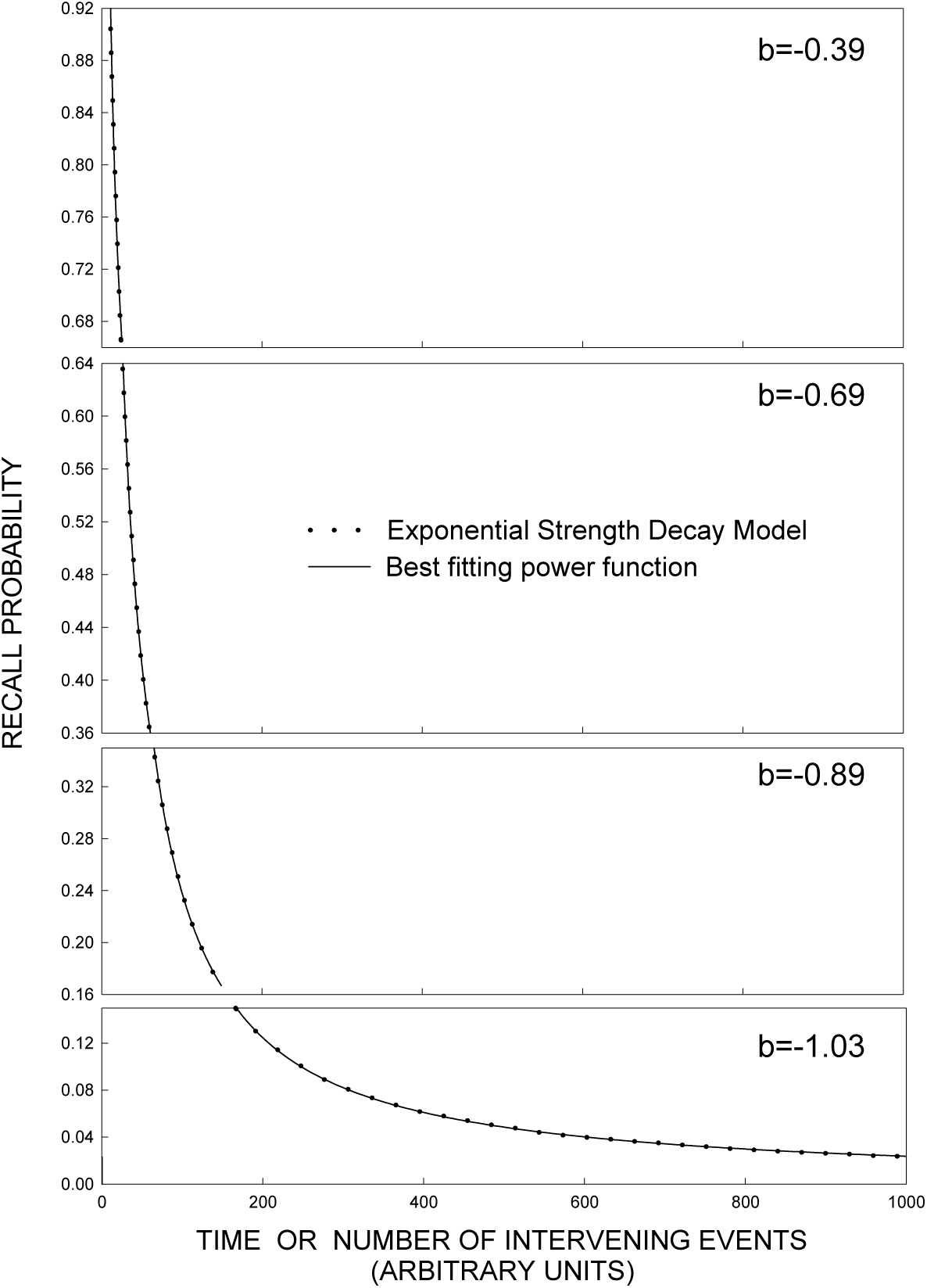
Replot of simulation example #3: A separate power function was fit to each of four regions of the forgetting function. The best fitting exponent, *b* is given in each panel.

Although our model assumes a constant threshold, it is mathematically indistinguishable from a model that assumes different thresholds for different items. To eliminate a free parameter, we allowed the variability in the *y*-intercept of the strength decay function forgetting function to subsume variability in the threshold (across items, but not time).

### 1.8. Conclusions

We have shown a striking example of how tricky it is to infer psychological process from the form of the retention function. Moreover, our analytic results suggest that there is something special about the power function as a description of the forgetting process. It may be that power functions arise from aggregate forgetting data under fairly general conditions. If this is true, then the challenge for memory theory is to explain violations of the power law of retention. One limitation of our analysis is that it does not readily handle RT data (this is because we assume a threshold that translates memory strength into binary responses). For learning data, the decrease in RT with practice obeys a power law (see Anderson, 1995 for a review). Regarding forgetting data, the increase in correct recognition RT with study-test lag is often linear (see Murdock, 1974, for a review) and the increase in inter-response times with output position is typically exponential (e.g., Murdock & Okada, 1970; Rohrer & Wixted, 1994; Wixted & Ebbesen, 1997). These findings are not addressed by our analysis.

In this note, we have shown how a simple model that assumes monotonic strength decay at the level of individual items coupled with a discrete retrieval threshold predicts aggregate power forgetting functions. This exploration illustrates some of the dangers inherent in inferring mental organization from the mathematical form of the behavioral data. The important question for theories of memory to address is how forgetting is affected by experimental manipulations and not what mathematical form the forgetting process assumes.

## ACKNOWLEDGMENTS

The authors acknowledge support from National Institute of Health grant MH55687 to Brandeis University. Correspondence concerning this article should be addressed to Michael Kahana, Volen National Center for Complex Systems, MS 013, Brandeis University, Waltham, MA 02254-9110.

## Footnotes

1 Consider the simulation shown in Figure 3. For these parameters, performance is at or near ceiling for small *t*. Although a power function accounts for nearly 100% of the variance when fit to the data from *t* = 1000 to *t* = 5000, the same power function does not provide a good fit when the data from *t* = 0 to *t* = 1000 are included.

## REFERENCES

Anderson, J. R. (1995). Learning and memory: An integrated approach. New York: Wiley.

Anderson, R. B. & Tweney, R. D. (1997). Artifactual power curves in forgetting. Memory & Cognition, 25, 724–730.

Anderson, R. B. & Tweney, R. D. (1998). The power law as an emergent property. Unpublished, Manuscript.

Bower, G. (1967). A multicomponent theory of the memory trace. In The Psychology of Learning and Motivation: Advances in Research and Theory, K. W. Spence & J. T. Spence, eds., vol. 1, New York: Academic Press, pp. 229–325.

Ebbinghaus, H. (1885/1913). Memory: A contribution to experimental psychology. New York: Teachers College, Columbia University.

Howard, M. W. & Kahana, M. J. (2002). A distributed representation of temporal context. Journal of Mathematical Psychology, 46.

Mensink, G.-J. M. & Raaijmakers, J. G. W. (1988). A model for interference and forgetting. Psychological Review, 95, 434–455.

Murdock, B. B. (1974). Human memory: Theory and data. Potomac, MD: Erlbaum.

Murdock, B. B. (1982). A theory for the storage and retrieval of item and associative information. Psychological Review, 89, 609–626.

Murdock, B. B. (1997). Context and mediators in a theory of distributed associative memory (TODAM2). Psychological Review, 104, 839–862.

Murdock, B. B. & Kahana, M. J. (1993). Analysis of the list strength effect. Journal of Experimental Psychology: Learning, Memory and Cognition, 19, 689–697.

Murdock, B. B. & Kahana M. J. (1993b). List-strength and list-length effects: Reply to Shiffrin, Ratcliff, Murnane, and Nobel. Journal of Experimental Psychology: Learning, Memory and Cognition, 19, 1450–1453.

Murdock, B. B. & Lamon, M. (1988). The replacement effect: Repeating some items while replacing others. Memory & Cognition, 16, 91–101.

Murdock, B. B. & Okada, R. (1970). Interresponse times in single-trial free recall. Journal of Verbal Learning and Verbal Behavior, 86, 263–267.

Rohrer, D. & Wixted, J. T. (1994). An analysis of latency and interresponse time in free recall. Memory & Cognition, 22, 511– 524.

Rubin, D. C. & Wenzel, A. E. (1996). One hundred years of forgetting: A quantitative description of retention. Psychological Review, 103, 734–760.

Wickens, T. D. (1998). On the form of the retention function: Comment on Rubin and Wenzel (1996): A quantitative description of retention. Psychological Review, 105, 379–386.

Wixted, J. T. (1990). Analyzing the empirical course of forgetting. Journal of Experimental Psychology: Learning, Memory & Cognition, 16, 927–935.

Wixted, J. T. & Ebbesen, E. B. (1991). On the form of forgetting. Psychological Science, 2, 409–415.

Wixted, J. T. & Ebbesen, E. B. (1997). Genuine power curves in forgetting: A quantitative analysis of individual subject forgetting functions. Memory & Cognition, 25, 731–739.

